# OPLS-based Multiclass Classification and Data-Driven Inter-Class Relationship Discovery

**DOI:** 10.1101/2024.09.23.614438

**Authors:** Edvin Forsgren, Benny Björkblom, Johan Trygg, Pär Jonsson

**Affiliations:** Computational Life Science Cluster (CLiC), Department of Chemistry, Umeå University, Umeå, Sweden; Department of Chemistry, Umeå University, Umeå, Sweden; Sartorius Corporate Research, Umeå, Sweden

## Abstract

Multiclass datasets and large-scale studies are increasingly common in omics sciences, drug discovery, and clinical research due to advancements in analytical platforms. Efficiently handling these datasets and discerning subtle differences across multiple classes remains a significant challenge.

In metabolomics, two-class OPLS-DA (Orthogonal Projection to Latent Structures Discriminant Analysis) models are widely used due to their strong discrimination capabilities and ability to provide interpretable information on class differences. However, these models face challenges in multiclass settings. A common solution is to transform the multiclass comparison into multiple two-class comparisons, which, while more effective than a global multiclass OPLS-DA model, unfortunately results in a manual, time-consuming model-building process with complicated interpretation.

Here, we introduce an extension of OPLS-DA for data-driven multiclass classification: Orthogonal Partial Least Squares-Hierarchical Discriminant Analysis (OPLS-HDA). OPLS-HDA integrates Hierarchical Cluster Analysis (HCA) with the OPLS-DA framework to create a decision tree, addressing multiclass classification challenges and providing intuitive visualization of inter-class relationships. To avoid overfitting and ensure reliable predictions, we use cross-validation during model building. Benchmark results show that OPLS-HDA performs competitively across diverse datasets compared to eight established methods.

This method represents a significant advancement, offering a powerful tool to dissect complex multiclass datasets. With its versatility, interpretability, and ease of use, OPLS-HDA is an efficient approach to multiclass data analysis applicable across various fields.

## Introduction

Partial least squares or Projection to Latent Structures (PLS) regression was first introduced in the 1970s by Herman Wold to handle cases where there are more descriptor variables than observations.^1^ Back then, PLS was built to predict a single response variable. Since then, it has evolved in several steps^2^ with Orthogonal PLS (OPLS) being introduced in 2002.^3^ OPLS simplifies interpretation by dividing the variation in descriptor variables into two parts: “predictive” variation related to the response(s) and “orthogonal” variation i.e. not related to the response(s). Regarding predictive power, OPLS and PLS are identical for single response variable models but due to the simplified interpretation, OPLS has gained popularity in fields where understanding the connection between descriptor- and response variables is crucial. Often, the response variable is categorical. Although regression was the original use of both PLS and OPLS, they are now commonly used for classification or Discriminant Analysis (DA) to distinguish between two or more groups or classes of samples. This type of classification works well for two classes, both in terms of discrimination and interpreting differences between classes. Several methods have been developed for interpreting differences between classes in two-class OPLS-DA models e.q. statistical total correlation spectroscopy (STOCSY),^4^ the S-plot^5^ and Selective-Ratio.^6^ The latter was initially developed for PLS-DA but later adopted for OPLS-DA.^7^ However, as the number of classes increases, OPLS-DA suffers in both interpretability and predictive power. The decline in performance is due to the “one-vs-rest” classification approach used in OPLS-DA models. This setup presents a challenge when one class is positioned between two others, which, combined with the linear nature of PLS and OPLS, results in weak models and complex interpretation.

A common approach to deal with this problem is to make multiple pairwise comparisons, “one-vs-one” models. These models can then be compared using SUS-plots (Shared and Unique Structure).^5^ SUS-plots are especially useful when multiple groups are compared against one control or reference group, such as different genotypes vs. wild-type or different treatments vs. untreated controls. This manual approach works well when the number of classes is small but quickly becomes challenging to manage as the number of classes increases. Despite these challenges, pairwise OPLS-DA is still widely adopted in omics for interpreting biological differences between different classes. Applications include differentiating brain cancer types,^8^ metabolomics in diet intervention studies,^9^ metabolite distribution patterns,^10^ and gene feature selection with microarray data.^11^ While manually creating multiple “one-vs-one” OPLS-DA models has been proven useful, it is a time-consuming process, both when creating the models and to summarize them into comprehensive results.

To circumvent most of the manual work, the Automatic Hierarchical Model Builder (AHiMBu)^12,13^ was introduced to handle these multiclass cases. And, although more effective than PLS-DA and a manual approach, there is still improvements to be made in terms of flexibility and interpretability.

Here, we introduce a data-driven methodology that combines two proven frameworks Hierarchical Cluster Analysis (HCA) with OPLS-DA, to create OPLS-Hierarchical Discriminant Analysis (OPLS-HDA). OPLS-HDA is developed to handle multiclass datasets where there are different levels of variation to reveal the hierarchical relations between classes whilst still detailing the discriminatory level. Employing OPLS-DA models as the foundation and a flexible approach to the hierarchical structure, provides OPLS-HDA with significant ad-vantages over existing methods. In addition to increased interpretability and flexibility, we also show that OPLS-HDA has competitive classification abilities compared to other popular methods on a wide range of datasets.

## Theory

### Orthogonal Projections to Latent Structure Discriminant Analysis - OPLS-DA

OPLS is a supervised multivariate projection method that uses latent variables to establish a linear relationship between the descriptor variables in the predictor matrix (**X**) and a response vector (**y**) or matrix (**Y**) for a specified set of observations. OPLS can be used even when the number of observations is smaller than the number of variables. It can handle correlated variables, noisy variables and missing values. This robust and versatile technique is applicable to a wide range of analytical tasks, including regression^3^ and discrimination,^14^ the latter being facilitated through OPLS-DA. In OPLS-DA, the response matrix (**Y**) is a dummy matrix that contains the information about class membership for each observation. OPLS separates the variation described by the model into two different parts, predictive and orthogonal. The predictive part is the variation in X (Eq. (1)) that is used to model the variation in **Y** (Eq. (2)). The orthogonal part contains variation in **X** (Eq. (1)) that is unrelated to the response **Y**. In the OPLS-DA context the predictive part contains between class variation while the orthogonal part contains within class variation. By dividing the variation into two part the interpretation of the model becomes easier.

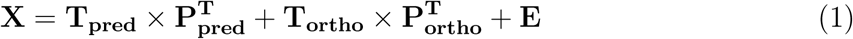

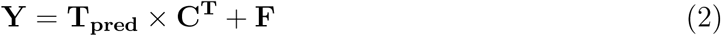

### Cross-Validation - CV

By dividing the training set into *k* groups and iterating over the groups, leaving one group out at a time, we can validate our model and get an estimation on how well it will perform on a test set. The cv-predictions i.e. prediction of samples when they are left out of the model denoted as *ŷ_cv_*, are used to determine the number of components to use in the final model by using the prediction error to decide whether to keep an additional component or not. The *ŷ_cv_* and the cv-scores of the samples in the training dataset are also useful to understand and interpret the data in a more realistic way than pure predictions of samples that are present in the model and is especially useful if data is scarce.^15^

### Distance Metric

The cv-predictions from two-class OPLS-DA models can be used to estimate the separability between classes. A suitable distance metrics for this is Cohen’s d, ^16^ defined in Eq. (3), which takes into account both the distance between classes and variation within them.

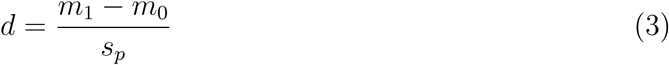

Here, *m*_0_ and *m*_1_ are the means of the *ŷ_cv_* of the two classes and *s_p_* the pooled standard deviations defined as in Eq. (4):

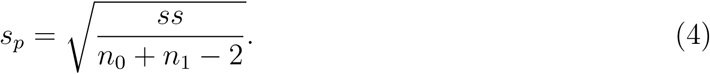

Here, *ss* is the pooled sum of squares of the two classes and *n* the number of observations in each class. The distance between the class means represents how well the OPLS-DA model can separate the classes, while the pooled standard deviation is a measure of the variability within the two classes. A model that can clearly distinguish between the two classes will result in a strong and valid model, characterized by predicted responses that are close to the actual targets (0 or 1) and a small pooled standard deviation, leading to a large Cohen’s d value. Conversely, in the case of weak models, the predictions often cluster around the value of 0.5 for samples from both classes. This clustering is typically accompanied by a large standard deviation, which indicates poor separation between the classes. Consequently, such a scenario results in a small Cohen’s d value.

### Hierarchical Cluster Analysis - HCA

Hierarchical Cluster Analysis (HCA) seeks the hierarchy of groups or individual observations based on a symmetrical distance matrix. There are two types of HCA, “bottom-up” (agglomerative) or “top-down” (divisive). For agglomerative, each observation starts as their own cluster and is then grouped successively. For divisive, all observations start in one cluster and are then divided to smaller ones successively. In this paper, we will refer to agglomerative HCA as HCA.

## Methodology

An OPLS-HDA model integrates the global interpretation and visual framework of dendrograms created with HCA with the predictive strength and detailed interpretation benefits of two-class OPLS-DA models. Put simply, an OPLS-HDA model is a top-down decision tree with a two-class OPLS-DA model in each decision node.

To create the hierarchy of the OPLS-HDA model, the relationships between all classes are mapped. This mapping is achieved by calculating all combinations of one-vs-one OPLS-DA models. Based on these models, the Cohen’s d between all the classes is calculated and stored in a distance matrix. This matrix is then used to create a dendrogram that forms the basis of the top-down decision tree populated with two-class OPLS-DA models. This integration creates a model that is interpretable at two levels: the hierarchical level, where the structure of the decision tree reveals the similarities and differences between classes, and the discrimination level, where the OPLS-DA models reveal details regarding how classes or groups of classes are differentiated from one another. An overview is shown in Fig. 1 and a detailed description of the setup and the model building is given below.

**Figure 1:**
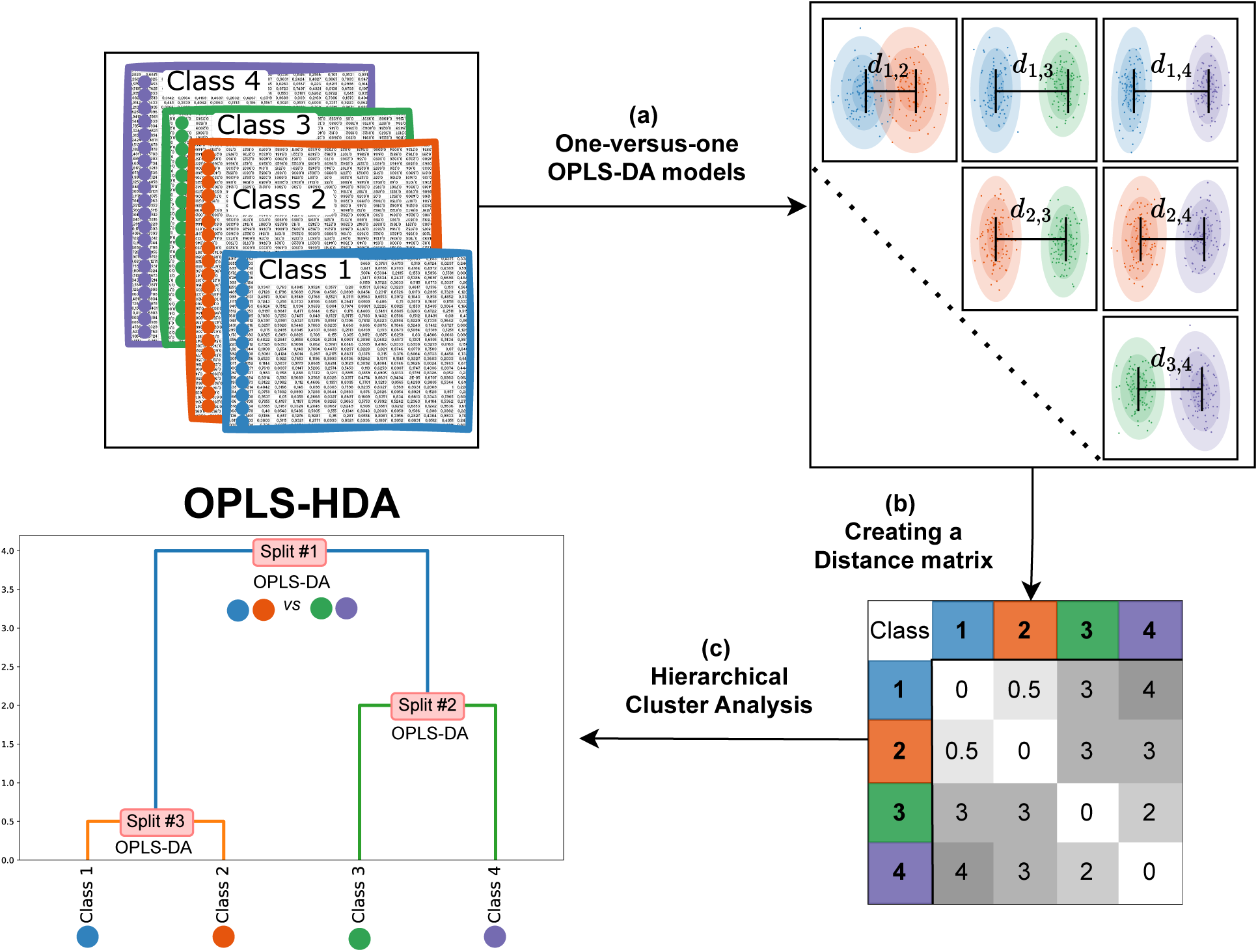
Observations of known classes, characterized by a common set of variables, serve as the starting point. The next step, (a), involves calculating all possible pairwise OPLS-DA models. From these models, Cohen’s d between classes is derived. These distances are compiled into a distance matrix, (b), which then forms the basis for hierarchical clustering analysis with a dendrogram as an end result, (c). The dendrogram not only reveals the relationships between classes but also functions as the basis of a decision tree, with a two-class OPLS-DA model at each split. This dendrogram, in conjunction with OPLS-DA models, constitutes the OPLS-HDA model.

The method is outlined in Fig. 1. All pairwise combinations of classes are compared using OPLS-DA models (Fig. 1a. The total number of models calculated in this step is 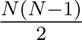, where *N* is the number of classes. This step is the most time-consuming part of the process, but when automated, still easily doable using a standard computer. The calculated distances are summarized in a symmetrical distance matrix, with each element representing the distance between two classes (Fig. 1b). The matrix captures the relational dynamics among all classes. The distance matrix is used to reveal the hierarchical level of the data in a dendrogram using HCA (Fig. 1c). The dendrogram is then populated with decision making OPLS-DA models in each split to create the decision tree, i.e, the OPLS-HDA model.

When using the OPLS-HDA model for classification, an observation begins at the top of the decision tree and moves downwards based on the predictions of each OPLS-DA model. It ultimately reaches one of the bottom leaves, indicating the class to which the observation belongs.

### Illustrative comparison of OPLS-HDA and OPLS-DA

To illustrate OPLS-HDA and show its benefits even in simple cases, we compared it to OPLS-DA using the Iris three-class dataset. ^17^ For simplicity in visualization, we limited the dataset to only two variables: petal length and petal width.

OPLS-DA inherently struggles with this relatively simple dataset since the decision boundary of an OPLS-DA model will always have a common center-point, due to its “one-vs-rest” setup. This causes trouble if we have more than two classes and the classes are separated along the same variables but at different scales. This is the case for the Iris dataset. When fitting an OPLS-DA model to a simplified version of this dataset the common center-point results in a suboptimal decision boundary (Fig. 2a). OPLS-HDA handles the placement of decision boundaries in a much better way (Fig 2b). By first calculating the distances between the three classes and performing HCA, we create a dendrogram (Fig. 2c), that acts as the foundation for the OPLS-HDA model. We then create OPLS-DA models in split #1 and #2 (Fig. 2d and e). The dendrogram complete with the two two-class models creates the decision tree and makes up the OPLS-HDA model. The resulting decision boundary of the OPLS-HDA model is a combination of the models in split #1 and #2 shown in Fig. 2b.

**Figure 2:**
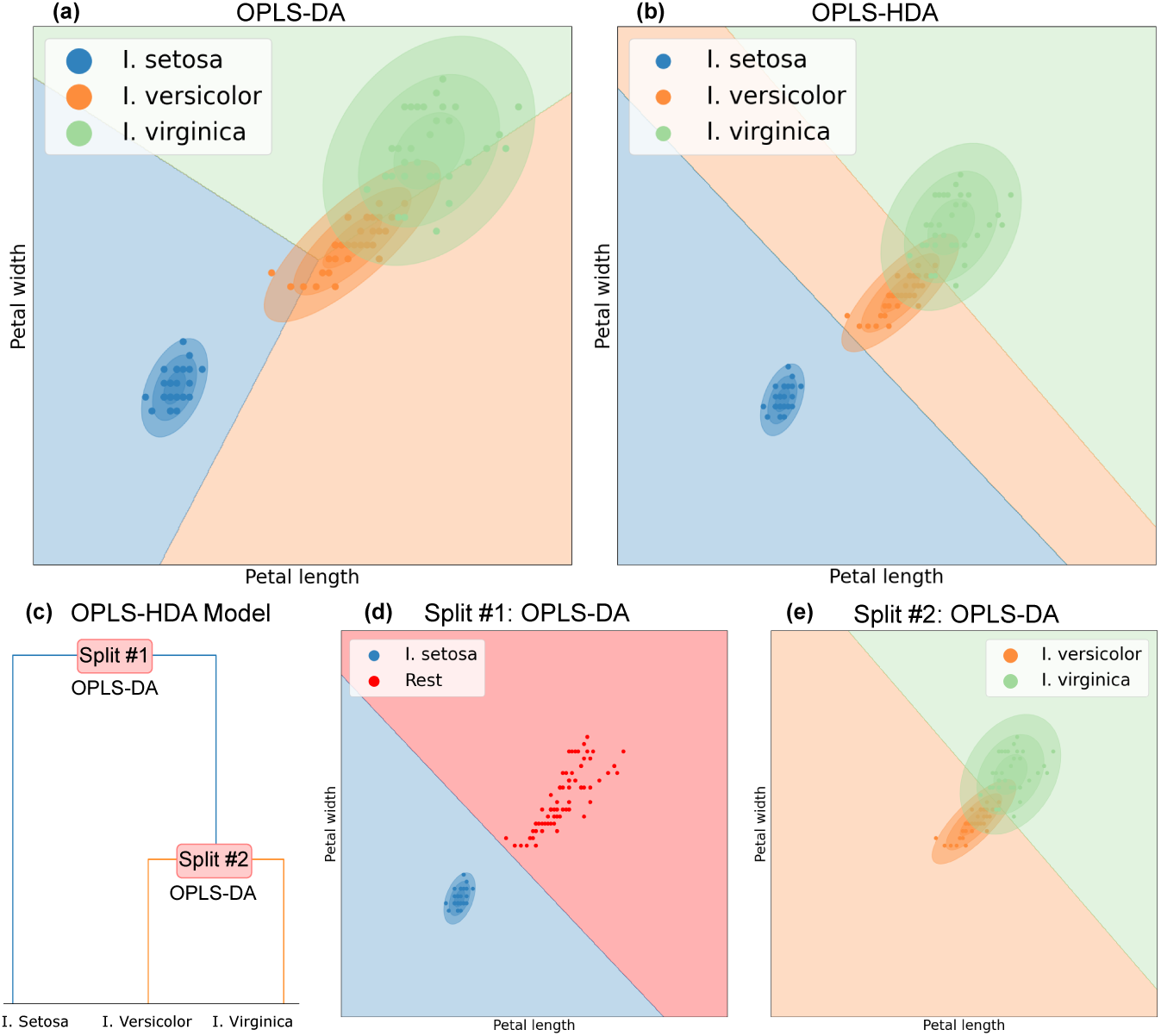
In (a) and (b) the decision boundaries of an OPLS-DA and an OPLS-HDA model are displayed. The OPLS-DA model has a common center-point leading to a sub-optimal decision boundary while OPLS-HDA creates a suitable boundary. In (c), the dendrogram on which the OPLS-HDA model is based. In each split of the dendrogram, a two-class OPLS-DA model is created. In (d), the decision boundary of the model in split #1 is shown and in (e), the decision boundary of the model in split #2.

### Software

For the OPLS-DA, we have utilized SIMCA, version 18.0.0 (Sartorius Stedim Data Analytics, Umeå, Sweden), as the backbone for model building including the cross-validation scheme. In HCA, the hierarchy can be created by linking observations differently by choosing a specific linkage algorithm. Throughout this work, we have applied SciPy to perform the HCA calculations with existing linkage choices. ^18^ The benchmarks presented in Supplementary information are performed in python using Scikit-learn^19^ and Eigenvector for AHiMBu.^12,13^

### Datasets

#### Glioma subtypes - Metabolomics

The brain tumor metabolomics data includes 232 observations and 240 identified metabolites originating from cross-platform GC-MS and LC-MS/MS global metabolomics analysis.^8^ The observations were subclassified based on molecular classifications according to WHO 2016 classification of tumors of the central nervous system. According to this molecular and histopathological classification, the material is distributed over seven adult glioma sub-types: 1. glioblastoma, IDH-wildtype; 2. glioblastoma, IDH-mutant; 3. astrocytoma, IDH-wildtype; 4. astrocytoma, IDH-mutant; 5. oligodendroglioma, IDH-mutant and 1p/19q-codeleted; 6. oligodendroglioma, NEC (IDH-wildtype and 1p/19q-codeleted); and 7. gliosarcoma, IDH-wildtype.

#### Whitefish - NIR Spectra

The Whitefish dataset includes 1,311 observations and 125 variables originating from Near-Infrared (NIR) Spectra data. The dataset contains 12 classes consisting of 4 fish species in 3 different states; frozen, fresh or thawed providing two main sources of class variation. This dataset is a robust use case for evaluating the efficacy of classification in scenarios where the variable space is correlated with a complex class structure. While the full dataset included 18 classes, we decided to restrict our study to a subset of 12 classes including species that had all three states measured.^20^

## Results

### Metabolomics Glioma - Streamlined identification of distinct metabolic characteristics

In the previous study, ^8^ a manual one-vs-one approach was performed to understand the relationship between the different classes. The findings where then summarized in a dendrogram with HCA and volcano plots were generated based on the clustering. This process involved a lot of manual work, the results of which we can efficiently replicate with OPLS-HDA.

To illustrate this, we applied OPLS-HDA to the high-dimensional mass spectrometry-based metabolomics data from the brain tumor tissues. The data included seven tumor subtypes, based on the WHO classification for tumors of the nervous system, including both molecular and histopathological analysis.^21^ Using tumor metabolic phenotypes as input, OPLS-HDA groups the seven glioma subtypes according to their WHO classification in a dendrogram (Fig. 3a). The dendrogram, uses Cohen’s d on the y-axis to reflect class proximity, providing an intuitive understanding of the relationships between the classes. The way OPLS-HDA visualize the data expedites initial data analysis, making it particularly useful of understanding major differences as well as unique features of defined classes in complex data, such as clinical omics data. Split #1 reveals a pronounced distinction between isocitrate dehydrogenase wild type (IDH-wt) and mutated tumors. This separation can also be visualized through traditional cv-score plots from the two-class OPLS-DA model (Fig. 3b).

**Figure 3:**
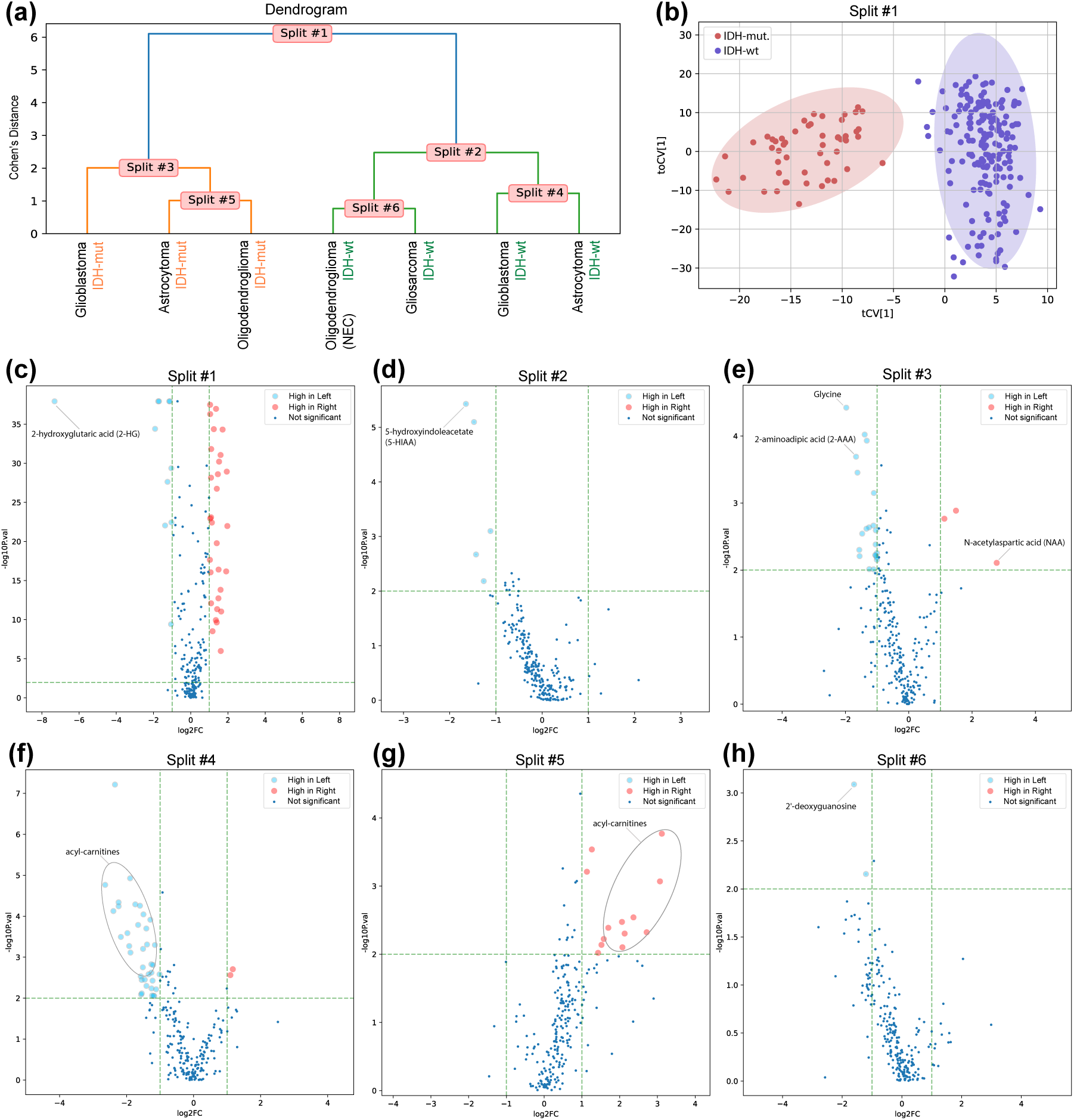
OPLS-HDA results of brain tumor metabolomic data. (a) Dendrogram for the brain tumor metabolomics data of seven WHO defined glioma subtypes. NEC = not elsewhere classified. (b) Cv-score plot for the two-class OPLS-DA model in (a), Split #1, separating IDH wildtype and IDH mutated tumors. (c)-(h) Volcano plots for each two-class OPLS-DA model in (a), Split #1-6. A few important metabolites, based on statistical significance (P*<*0.01) and expression level (*>*2-fold difference), are highlighted. Metabolite fold differences are shown as ratios according to the analyzed Split, as illustrated in the dendrogram in (a). I.e “High in Left” or “High in Right” refers to left-or right-hand side of the Split shown in (a).

To further investigate the unique features driving the separation of the different tumor classes, we generate volcano plots for each two-class OPLS-DA model, based on statistical significance and expression level for individual metabolites (Fig. 3c-h). This analysis shows that Split #1, which separates the three IDH-mutated classes from the four IDH-wt classes, is driven by 41 metabolites with significantly different levels in the tissues (P*<*0.01 and *>*2-fold difference). The most important metabolite is the high expression of 2-hydroxyglutaric acid (2-HG) produced in IDH-mutated cells (Fig. 3c).

Focusing on the subsequent splits, we see that Split #2 differentiates IDH-wt classes; glioblastoma and astrocytoma from oligodendroglioma (NEC) and gliosarcoma. This differentiation is primary driven by high levels of 5-hydroxyindoleacetate, a metabolite of serotonin catabolism (Fig. 3d).^8^ Turning our attention to the IDH-mutated classes, Split #3 distinguishes high-grade glioblastoma from lower-grade astrocytoma and oligodendroglioma tumors. Metabolic markers for rapid cell proliferation, such as glycine and 2-aminoadipic acid, were prominent in high-grade glioblastoma while N-acetylaspartic acid, a metabolic marker for normal nervous tissue, was low as expected (Fig. 3e). As we move down the dendrogram, the differences between analyzed classes becomes less pronounced, which is anticipated as the tumors metabolic phenotypes becomes more similar. However, both Split #4 and Split #5, regardless of IDH mutation status, show reduced levels of a broad range of acyl-carnitines in lower-grade astrocytoma tumors, highlighting these metabolites as unique features (Fig. 3f-g).

### Whitefish dataset - Hierarchy and score plots

In the Whitefish study, 66 one-vs-one OPLS-DA models are summarized and presented in a dendrogram, as shown in Figure 4a. This dendrogram provides clear insights of how different groups relate to each other, namely that the state of the fish is the main source of variation and thus frozen samples differ significantly from fresh or thawed ones. Speculatively, this is because of the presence of ice in the frozen samples. Next, we see that the same species that are either fresh or thawed are more similar to each other and that cod stands out distinctly from other species, regardless of whether the samples are frozen or fresh/thawed.

**Figure 4:**
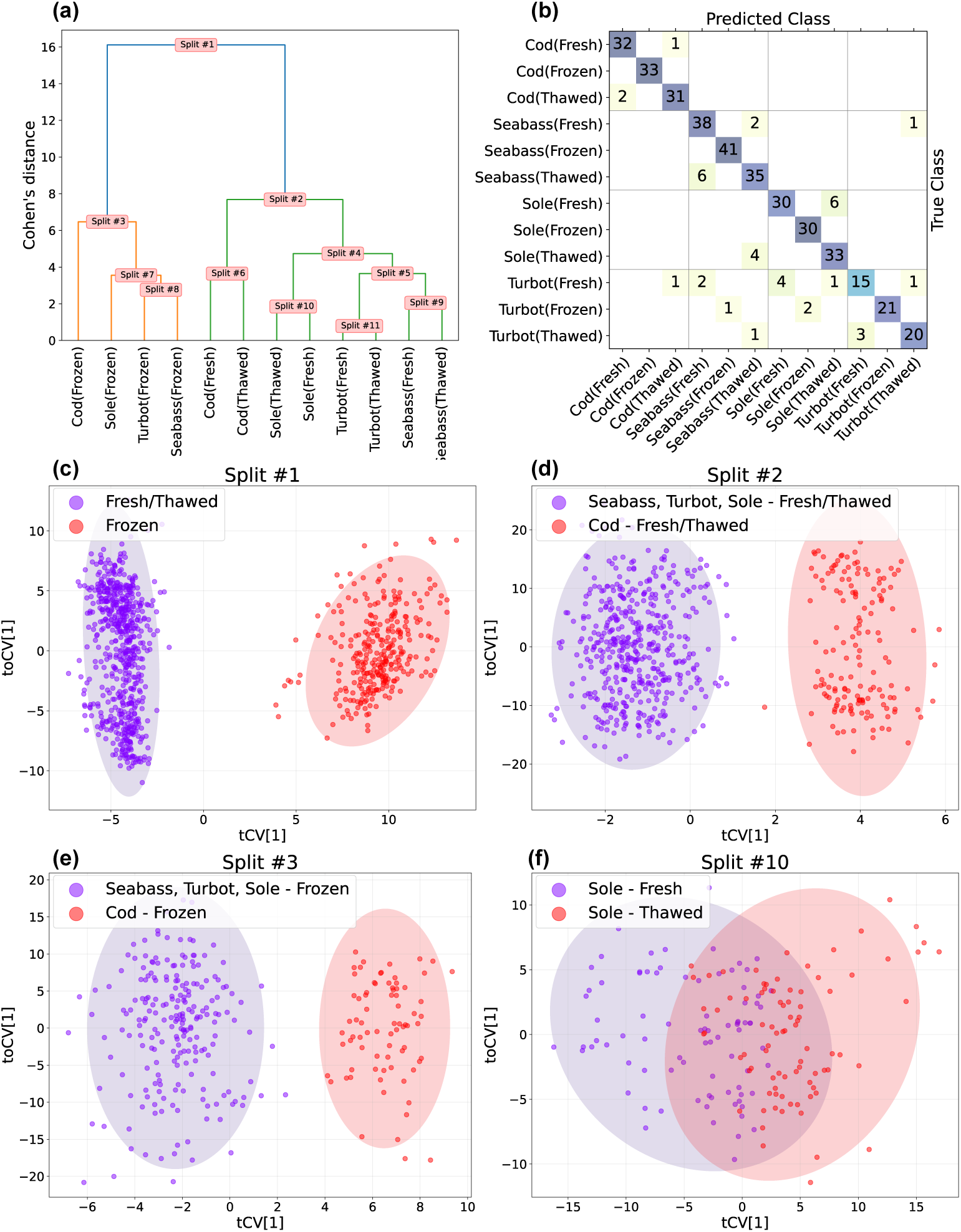
(a) Dendrogram of the whitefish data produced by the Cohen’s d calculated from the *ŷ_cv_* of 66 one-vs-one OPLS-DA models. Frozen classes are separated from fresh/thawed and cod is separated from the other species. (b) Confusion matrix of the test set with an accuracy of 90.4% with the actual class belongings on the y-axis and the predicted on the x-axis. (c-f) Cross-validated score plots from two-class OPLS-DA models in splits #1, #2, #3 and #10, from the OPLS-HDA model shown in (a). Split #1-3(c-e) are clearly separated while #10 has a significant overlap. The ellipses around each class in the score-plots (c,d,e & f) are 95% confidence intervals for each class based upon Hotelling’s T2.

These observations are supported by the cv-score plots from the OPLS-DA models in the splits, shown in Figures 4c-f. Similarly, the confusion matrix in Figure 4b confirms these find-ings i.e. as we move down the tree, the likelihood of correctly classifying a sample decreases. Notably, no frozen sample was misclassified as fresh or thawed. Also, cod is consistently classified separately from other species. The primary challenge involves confusing fresh and thawed samples within the same species, as illustrated in Figure 4f, where the cross-validated scores for fresh and thawed sole overlap significantly.

### Benchmark results

To compare OPLS-HDA with other multiclass classification methods, we have performed a benchmark on a wide range of datasets. The datasets range from simple ones as Iris ^17^ to high-dimensional once such as the Breast Cancer dataset.^22^ We compare the performance of OPLS-HDA to eight other methods (Table 1). From this, OPLS-HDA emerges as a strong and flexible competitor to the other methods. Being in the top three for all datasets except LIVECell results in OPLS-HDA having the highest average accuracy of all the tested methods. This displays the robustness and flexibility of OPLS-HDA to handle both low- and high-dimensional datasets with varying characteristics. A more in-depth description of the datasets and the tested methods are presented in Supplementary materials.

**Table 1:**
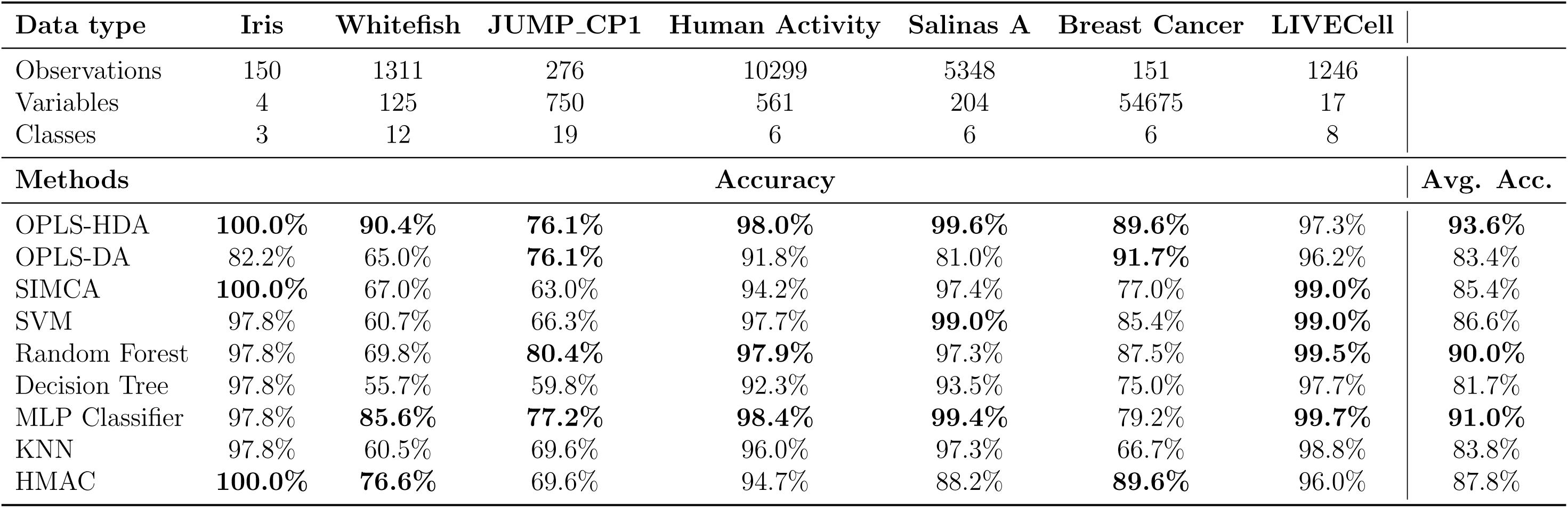
Combined dataset summary and benchmarking results. The top-three methods are highlighted in bold.

## Conclusion

In this paper, we have introduced a new method for multiclass classification called Orthogonal Partial Least Squares-Hierarchical Discriminant Analysis (OPLS-HDA). This method is based on OPLS, a widely used chemometric tool with applications in both industry and academia. The motivation behind this method is that OPLS-Discriminant Analysis (DA), which can be used for multiclass classification, often struggles as the number of classes increases. For two-class classification, OPLS-DA is highly effective, offering strong discrimination between classes and providing interpretable information on why the two classes differ. OPLS-HDA integrates the strengths of two-class OPLS-DA models with HCA to create a top-down OPLS-DA-based decision tree.

In the glioma study, we showed that OPLS-HDA results in much less manual work but with the same interpretation as the original paper.^8^ The whitefish NIR dataset differ in both state and species where OPLS-HDA clearly separated the biggest source of variation to smaller sources. Further, we have benchmarked OPLS-HDA on eight datasets and compared the classification test set accuracy to eight popular machine learning models.

We demonstrate that with OPLS-HDA’s data-driven decision-making, we avoid tedious manual work while still allowing for a detailed interpretation of the difference between classes. This approach allows researchers to focus on interpreting their data and find the answers to their research questions instead of a time consuming model-building process. This methodology not only streamlines the analytical process but also enhances the robustness and clarity of the results, paving the way for more efficient and insightful multiclass-research in the future.

## Acknowledgement

The authors thank Lennart Eriksson for reviewing and commenting on the text in the manuscript. The authors also thank Sean Roginski for his helpfulness and insights regarding our benchmark tests of AHiMBu.

## Supporting Information Available

### Benchmarking

To ensure OPLS-HDA is a robust multiclass alternative, we have performed a benchmark on a wide range of datasets. The datasets range from simple ones as Iris with 3 classes and 4 variables to truly high-dimensional once with tens of thousands of variables with a lot fewer observations. We compare the performance of OPLS-HDA to 7 other established methods.

#### Method descriptions

##### Soft Independent Modeling of Class Analogy - SIMCA

Soft independent modeling of class analogy (SIMCA) is a multivariate method where one PCA model is created for each class. To classify an observation, the residual, Distance to model in X-space (DmodX), is used to see if the observations fit into the PCA model or not. SIMCA is often used in process control to make sure that a process is not deviating from historical data. It is effective in these circumstances, but is a lot harder to interpret than OPLS-DA.^23–25^

##### Automatic Hierarchical Classification Model Builder (AHIMBU) - Hierarchical Model Automatic Classifier (HMAC)

Automatic Hierarchical Classification Model Builder (AHIMBU) or Hierarchical Model Automatic Classifier (HMAC) as the method is called in Eigenvectors implementation. AHIMBU is a method that tries to solve the same problems as OPLS-HDA with PLS-DA models. Similarly to OPLS-HDA, AHIMBU begins with creating one-vs-one models to map the relationship between classes. But then, the two classes that have the lowest success rate (highest classification error) are selected and merged into one class. The minimum cross-validated nonerror rate is used to estimate the classification error. If all classes then can be separated perfectly in one model, or if there are only two classes remaining, all the relevant models are built. If this is not the case, then the process is repeated. This will, similarly to OPLS-HDA, result in a decision tree based on PLS-DA models. However, OPLS-HDA can handle orthogonal variation, and utilizes Hierarchical Cluster Analysis (HCA) based on a distance matrix of statistical measures of effect (Cohen’s distance) to achieve its structure whereas AHIMBU relies solely on classification error.^12,13^ The difference becomes clear when the two methods are benchmarked on several datasets.

##### k-Nearest Neighbor

The k-Nearest Neighbors (k-NN) algorithm is a non-parametric method meaning that there is actually no “model” that needs to be trained or tuned. It instead classifies a data point by analyzing the ‘k’ closest labeled data points and adopting the most common class among them, based on a distance metric. The distance calculations for prediction are computationally intensive with big datasets but often works well as long as the space is not too high-dimensional.^26,27^

##### Decision Tree

The Decision Tree is a predictive method used in statistics, data mining, and machine learning. It uses a tree-like model of decisions and their possible consequences. Decision Trees are simple to understand and interpret, but they are prone to overfitting, and often perform poorly on test data.^28^

##### Random Forest

Random Forest is an ensemble learning technique constructed of multiple decision trees. The prediction is then based on a majority voting of the decision trees in the forest. This method addresses one of the key limitations of a single decision tree, which is the tendency to overfit the training data. A Random Forest can handle large datasets efficiently and is used in diverse applications such as credit scoring and disease prediction. However, for complex data, the understanding and interpretation of the decisions and predictions are challenging. ^29^

##### Suport Vector Machines

Support Vector Machines (SVM) is a popular method for classification that is especially efficient in high-dimensional spaces. SVM works by constructing a hyperplane or a set of hyperplanes in a high-dimensional space. This space can then be used for classification or regression. In classification, the core idea is to maximize the margin between the data points of different classes, effectively creating the widest possible gap between classes. However, tuning hyperparameters can be challenging to find an appropriate fit. Interpreting the decision boundary can be done in lower dimensions, but with growing dimensions the interpretability becomes challening.^30^

##### Multi-Layer Perceptron (MLP) - Artificial Neural Network (ANN)

Multi-Layer Perceptron (MLP) is a class of feedforward artificial neural network (ANN). An MLP consists of at least three layers of nodes: an input layer, a hidden layer, and an output layer. Except for the input nodes, each node is a neuron that uses a nonlinear activation function. MLP utilizes backpropagation to optimize the model. MLPs are widely applied in many fields but require large datasets and careful tuning of hyperparameters. As widely recognized, the interpretation of ANNs is hard.^31^

Each of these methods has its strengths and weaknesses, making them suitable for different types of problems in the field of machine learning. The choice of algorithm often depends on the size of the dataset, the nature of the problem, the computational resources available, and how valued interpretation.

### Datasets

The datasets were chosen to represent a wide range of different data types, number of classes, observations, and variables. This, to ensure OPLS-HDA is not only applicable on certain datatypes but performs well in most settings.

#### Iris

The Iris dataset spanns 150 observations across 3 classes. The variables are the lengths and widths of the sepals and petals of three species of iris plants. Although with only 4 variables, this dataset is widely used as a comprehensive benchmark example.^17^

#### Subset of CP1 JUMP

The subset of CP1 JUMP dataset, comprising 276 observations and 750 variables derived from CellProfiler features, and spans 19 classes of compound treatments. In this high-dimensional dataset, the variable space outweighs the observational space. It offers a challenging task for methods to effectively discern between numerous classes, all while combating the challenges posed by the curse of dimensionality.

#### Human Activity

The Human Activity dataset houses data from 10,299 observations, each described through 561 variables, and consists of 6 activity classes. The variables are drawn from cell phone accelerometers and describes patterns and trends in human motions. ^32^

#### Salinas A

In the domain of remote sensing, the Salinas A dataset, with 5,348 observations and 204 variables spread across 6 classes, is derived from hyperspectral images of land. The spectral data, challenges the methods with highly correlated variables.^33^

#### Breast Cancer

The Breast Cancer dataset, hosting data from 151 observations across 6 classes, is characterized by a variable space of 54,675 gene expression variables. It is a truly high-dimensional biological data scenario, with a very limited number of observations.^22^

#### LIVECell - Cell Morphology

The LICVECell dataset is a collection of phase-contrast images of cells from 8 different cell lines imaged at different time points.^34^ The cells are grown in wells and each well is imaged at two regions of interest (ROI). 17 image features (variables) describing each cell’s morphology have been extracted using single-cell data.For each region, the median of all cells is used, resulting in 1246 observations, split into a training and a test set. Each well could only be part of either the training or the test set to reduce overfitting and maximize the variability between the two sets of data.

These public datasets, each with their unique challenges and characteristics, provide a comprehensive and diverse array to validate the OPLS-HDA methodology across various domains, dimensionalities, and class structures. Consequently, they offer a wide view of its performance and applicability in varied real-world scenarios, ranging from simple and direct classifications to high-dimensional, multiclass situations.

